# A comparison of rotation-based scores for gene set analysis

**DOI:** 10.1101/2021.03.23.436604

**Authors:** Adria Caballe Mestres, Antonio Berenguer Llergo, Camille Stephan-Otto Attolini

## Abstract

Gene-wise differential expression is usually the first major step in the statistical analysis of high-throughput data obtained from techniques such as microarrays or RNA-sequencing. The analysis at gene level is often complemented by the screening of the data in a broader biological context that considers as unit of analysis meaningful groups of genes that may have functions in certain biological processes.

Among the vast number of publications about gene set analysis, the rotation test for gene set analysis, also referred by roast, is a general sample randomization approach that maintains the integrity of the intra-gene set correlation structure in defining the null distribution of the test.

In this work we compare the performance of several enrichment score functions using such rotational approach for hypothesis testing. We find that computationally intensive measures based on Kolmogorov-Smirnov statistics fail to improve the rates of simpler measures of GSA like mean and maxmean scores. We also show the importance of accounting for the gene linear dependence structure of the testing set, which it is linked to the loss of effective signature size. In this regard, weighted statistics are introduced with the aim of maximizing the effective signature size. These are found to out-power other usual scores in some simulations scenarios. The average of absolute values is found to be the most powerful score using both simulated and benchmarking data.

All tools are available in the **roastgsa** R package.

## 1 Introduction

Gene-wise differential expression is the most common step in the statistical analysis of high-throughput data generated with microarrays or RNA-sequencing. Subsequent analyses include the screening of the data at broader scales whose measurement unit are groups of genes with common biological functions.

There are a multitude of methods to evaluate the aggregated gene expression changes in functional gene sets under different experimental conditions. These are typically classified according to the statistical test [5]: (a) Self-contained approaches assess whether the observed gene set association to the experimental condition is expected by chance [11, 14, 9]; and, (b) Competitive approaches aim to determine if the observed association is expected in comparable gene sets [1, 19, 3, 22, 19, 7, 12, 23, 13].

The randomness associated to the underlying null hypothesis has been approximated by non-parametric approaches that use sample randomization [11, 14] - typically associated to self-contained testing-, gene randomization [19, 18] -competitive testing- and a hybrid of the two [19, 3], -also competitive testing-. Similarly, parametric approximations of either types have been developed in the past [7, 12, 22, 9].

Gene Set Enrichment Analysis (GSEA) [19] is one of the most widely used approaches for enrichment in the biomedical community. It computes a Kolmogorov-Smirnov (KS) test by comparing target gene sets with the rest of the genes in the genome. For a sufficiently large number of observations, sample permutations are used to maintain the integrity of the intra-gene set correlation structure in defining the null distribution of the test, resulting in a gene and sample randomization hybrid. However, for small sample sizes (less than 7 per experimental conditions), p-value granularity becomes a severe problem and it is recommended to apply permutation (or randomization) of genes instead. Nevertheless, this approach ignores the underlying correlation structure of the testing-set, leading to a higher false positive rate when the intra-gene set correlation is larger than expected in a randomly selected gene set [20]. Besides, the genomic profile of genes not in the testing set is used twice, first to calculate the test statistic and second to find the null distribution.

Smyth et al. [11] suggest the more general procedure of rotating the residual space of the data, which is useful even for small numbers of degrees of freedom. Both the self-contained test (roast) and its competitive version (romer) have been implemented [17]. The romer methodology can be considered as the most general gene and sample randomization GSA approach in the current literature [15] and it is the focus of attention of this work. Nevertheless, to our understanding, the test statistics provided in romer, which are all functions of the moderated t-statistics ranks, are too limited.

Frequently, gene set data bases such as KEGG, Hallmark or GO terms are used for battery testing. Some of the defined sets are composed by genes that interact in any particular biological process, leading to high intra-gene set correlations. Focusing on unadjusted GSEA figures in which a good part of the genes go to the same direction might be misleading to researchers as it can be hard to differentiate between true pathway activity and additive noise effect due to gene-to-gene dependence.

In this paper we review the rotational approach for linear models presented in [8], which motivates the roast method for enrichment, and pro-pose to complete the romer functionality by providing other statistics used in the GSA context. We compare the performance of the KS based test statistics introduced in the GSEA [19] and GSVA [6] methodology as well as restandardized statistics based on summary statistics measures [3] using both simulated and benchmarking data [21].

Furthermore, to interpret the output of the roast GSA methods, we encourage the usage of effective signatures size, a proxy for the number of independent genes in the testing gene set that can be directly linked to the power of the statistical test being used.

All investigated measures are implemented into the R package **roastgsa** [shortly in bioconductor].

## 2 Methods

### 2.1 Data sets

#### Metabric data

The metabric database contains the genomic profile and clinical information of 1980 patients with breast cancer [2]. We select the individuals based on their ER status (that takes state positive and negative). Only ER positive samples are selected. The linear gene dependence structure of the metabric data is used for simulations.

#### Gene set collections

The Hallmark gene set collection from MSigDB [10] is used to assess the performance of the gene set analysis methods presented in this paper. We consider the interferon alpha hallmark as motivation for our proposed methodology since it shows the highest intra-gene set correlation of all 50 evaluated gene sets. Both TNFA signaling via NFKB and interferon alpha, with medium and high intra-gene set correlations [Supp. Figure 1], respectively, are used for testing under simulated data.

#### Benchmarking data

A compendium of 42 microarray datasets investigating the KEGG specific functionality of human disease are compiled by [21] and made accessible through the R package GSEABenchmarkeR [4]. These benchmarking data are used to compare the performance of the presented GSA scores.

### 2.2 Rotations based gene set analysis

Rotation tests for multivariate linear regression were first proposed in [8] as a generalization of standard permutation tests, with the assumption of multinormality. If such distributional assumption is feasible, the rotation tests have the great advantage that can be used for complex models even when the sample size is very small. This makes the approach very appealing to deal with high-throughput data, where typically the number of samples available is limited, and hence p-value granularity is presented as a severe issue for permutation based approaches. In [11] the rotation approach is adapted so that can be used as the most general gene set enrichment analysis tool, both for competitive and self-contained testing.

Let *Y_*i*_* be a *q*-dimensional vector, independent for any *i* ∈ [1,…, *n*], that represents the gene expression profile of the *i*th individual with the following multivariate normal distribution assumption

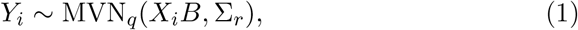

where *X* is a *n* × *p* design matrix with *p* – *k* adjusting covariates (the first *p* – *k*) and *k* covariates of interest. The *p* × *q* matrix *B* contains the linear regression coefficients and Σ_*r*_ is the error covariance matrix. The main steps of the rotational approach proposed in [11] can be summarized by:

1. QR decomposition of *X* to estimate the regression coefficients of interest for the *q* genes and their corresponding error variance.
2. When *q* is sufficiently large, moderated t, as defined in the **limma** methodology, can also be computed and used for further calculations of the enrichment score. The moderated t-statistic updates the error variance of the linear models using the information of all the estimated variances based on empirical Bayes posterior means. The prior distribution is obtained by fitting a scaled F-distribution to the sample variances. The posterior distribution is the weighted average of the estimated location of the prior distribution and the sample variances. Weights are determined by the degrees of freedom of the estimated F-distribution and *n* – *p*, respectively. Moderated-t statistics are further transformed to z-scores by using the quantile function of the Student-t distribution. This is especially useful when the number of degrees of freedom left in the model is small, and the observed t distribution is heavy-tailed.
3. For any testing gene sets, a GSA test statistic score is calculated using the moderated t’s. Depending on the proposed statistic, hypothesis testing will be considered either competitive or self-contained.
4. Rotation applied to the residual space of the data can be handled by conditioning only on sufficient statistics of the unknown covariance matrix Σ_*XY*_. Rotation statistics can be estimated and used to define the null hypothesis for hypothesis testing.

### 2.3 Defining the null hypothesis and GSA test statistics

Let [*δ*_*i*_, *i* ∈ Ω = [1,…,*q*]] be the *q*-vector with the moderated t-statistics, let *S* ⊂ Ω be the index vector with the position on [*δ*_*i*_] of the *m*_*o*_ = |*S*| genes in the set of interest with complement set *C* = Ω \ *S*, and let [*ρ*_*i*_] be the rank in decreasing order of the gene in position *i*.

In assessing the statistical relevance of a gene set of interest *S*, the first main consideration is defining whether the hypothesis testing is presented as a self-contained (SC) or a competitive (CO) problem. The self contained hypothesis compares the coefficients of the model for the gene set of interest against what could be found by chance if new data were observed, whereas the competitive approach makes the comparison taking into account the rest of the genome. Two null hypothesis assumptions can be placed in both type of testing problems: (distributional) equality in the distributions of the testing set 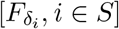 and the complementary set 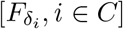, if competitive testing, or [*δ*_*i*_ = 0, ∀*i* ∈ *S*], if self-contained testing; (locational) equality in the first moment of the testing set [*δ*_*i*_, *i* ∈ *S*] and the complementary set [*δ*_*i*_, *i* ∈ *C*], say avg_*S*_ (*E*[*δ*_*i*_]) = avg_*C*_ (*E*[*δ*_*i*_]), if competitive testing, or avg_*S*_ (*E*[*δ*_*i*_]) = 0 if self-contained testing.

In Table 1 we present several GSA test statistics that can be used for biological enrichment with different goals and interpretations. The *T*_*mean*_ (both for CO and SC), *T*_*meanrank*_ (only CO) and *T*_*median*_ (CO and SC) maintain the integrity of both distributional and locational hypothesis. The mean, which we present as a weighted mean for its more general form, is provided to measure the common directional behaviour of the testing set. The median and meanrank are robust measures to outliers, that can prevent giving importance to gene sets with only few influential genes at the expense of losing statistical power. These scores can represent a suitable way to rank gene sets in battery testing when extreme values are undesirable. We also present the *T*_*maxmean*_ (CO and SC), the *T*_*ksmax*_ (CO) and the *T*_*absmean*_ (CO and SC). These three scores violate the locational null hypothesis unless the more restrictive distributional hypothesis is imposed. The maxmean, defined in its general form, uses the moderated-t magnitudes of only the most prominent direction, either positive or negative. This is relevant to pick up the main trend of the gene set without compromising the statistical power. The ksmax is the original score for GSEA [19]. Similarly to maxmean, it looks for concentration of the genes in the testing set in either one of the two extremes of the ranked list of genes (positive or negative). The absmean is the only undirectional score presented here which is found useful as a way to capture activity of genes in the testing set, regardless of their direction. Finally, the ksmean (CO) uses a similar KS statistic to the ksmax, but looks for a global effort of the genes in the testing set, that penalizes effects with contrary directions, hence it controls the rejection rate under the locational null hypothesis when the distributions in the two directions are equals.

**Table 1:**
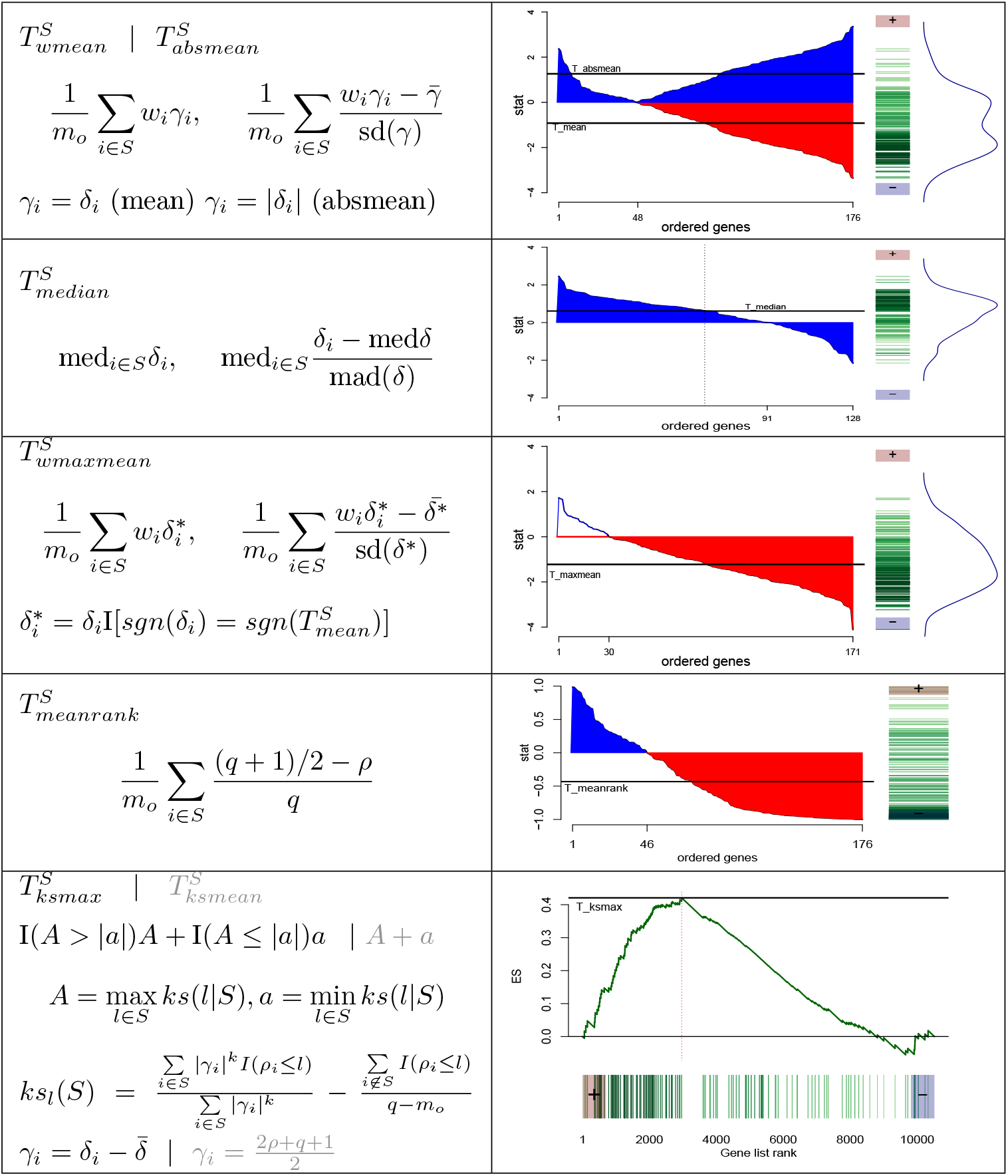
GSA Scores: The functions mean, maxmean, median and absmean are formulated for both SC (left) and CO (right) testing. The meanrank, ksmax and ksmean are exclusive scores for the competitive approach. The mean and maxmean approaches are expressed in a more general form that weights the gene importance in the score calculation [online methods 4].

### 2.4 Effective signature size of a gene set

Important pathways in database collections such as Broad Hallmarks or KEGG might contain highly correlated genes, even when the effect of the known covariates is adjusted *a priori* [Figure 1a shows the correlation structure of the Interferon alpha gene set using the metabric data]. This alters heavily the precision in estimating the gene set activity as, generally, the variance of the summary statistics increases with the intra-gene set correlation. As an example, for the mean GSA statistic presented in Table 1, assuming equal variances *σ*^2^ for all moderated t-statistics [*δ*_*j*_], for *j* ∈ [1,…,*m*_0_], it follows that var(*T*_*mean*_(*S*)) = *σ*^2^/*m*_0_ (1 + *ρ* (*m*_0_ – 1)) increases linearly with the average correlation *ρ*.

**Figure 1:**
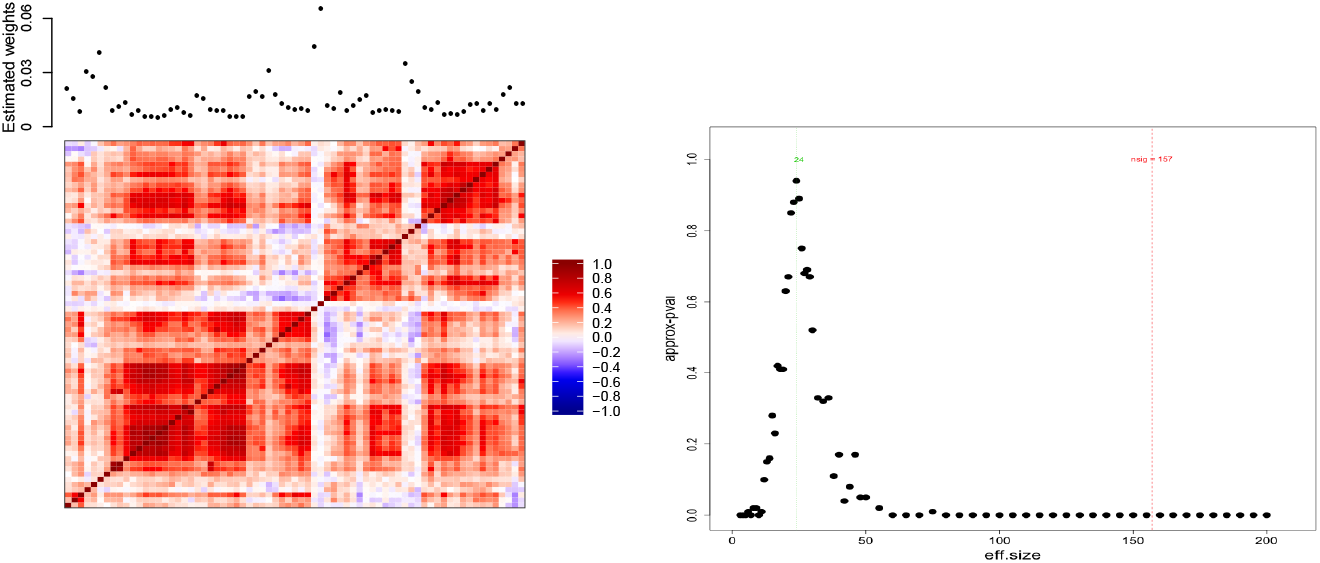
[Left] Heatmap with the interferon alpha gene-gene Pearson correlations (from −1 in dark blue to 1 in dark red). Estimated weights are shown above the heatmap. [Right] Effective signature size for the myogenesis hallmark as measured by the number of randomly selected genes to obtain similar levels of variability in the rotation scores.

We define the effective signature size of a given gene set *S* with size *m*_0_ and test statistic variance var(*T*_*any*_(*S*)) by the minimum number of independent random genes *m*_*e*_ needed to obtain test statistic variances as high as var(*T_any_*(*S*)). In this regard, we consider an approximation of the effective signature size, which we find more appropriate for our problem, by the number of randomly selected (not necessarily independent) genes that are needed to achieve comparable variability levels on rotation summary test statistics. For any pathway of interest, we compare the variance of the rotation scores of such tested set, say *υ*_*s*_, and the same variances for sets composed by randomly selected genes of size *m*, say 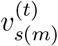, for any instance *t* ∈ [1,…, *T*]. We compute a p-value for the specific effective signature size *m* by

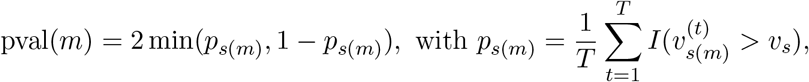

that approximates the probability of obtaining a variance as extreme as *v_s_* in a randomly selected set of size *m*.

In Figure 1b we present the approximated p-value curve as function of the effective signature size in a real case study (downsampling to *n* = 20 the metabric data and using the Myogenesis hallmark as testing gene set). We observe that random gene sets with less than 50 genes obtain similar levels of variability as the Myogenesis gene set, which defines a gene set of 157 genes.

### 2.5 Weighting summaries

We present a strategy to weight the gene importance in summary statistics for GSA to lower down the variance of the rotation scores, thus keeping a larger effective signature size [see Supp. 1 for further motivation to use weighted summaries].

To do so, we propose to weight down the importance of genes that are already highly explained by other genes in the testing gene set. For every gene in the gene set, we measure the variance explained by all other genes using a linear regression model. Since the interest here is not in the variable selection part of the regression problem, we consider a regression model only using the leading PCA scores (of the data ignoring the gene whose weights wants to be estimated) as linear predictors. PCA based regression is commonly used in high-dimensional data to (1) reduce the number of predictors and make the problem treatable (2) avoid collinearity problems (3) avoid over-fitting problems. Below, details to estimate the weights are provided.

#### Estimated weights

The signature size, denoted by *m*_*o*_, defines the number of (typically) genes that compose the pathway of interest, say *S*, that wants to be tested. If *w*_*i*_ = 1/*m*_*o*_, for all *i* ∈ *S*, then the statistics coincide with their unweighted versions. We propose to calculate the weights as follows. Let *P*^(*g*)^ be the *n*×*k* matrix with the scores of the first *k* principal components applied to the data without the *g* row, [*Y*]_*S*\*g*_, for all *g* ∈ *S*. Then 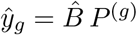 is the fitted values of the linear model that associates *Y*_*g*_ (dependent variable) and *P*^(*g*)^ (independent variables), with degrees of freedom *df* . The weights are calculated based on the coefficient of determination of all *m*_0_ fitted linear models by

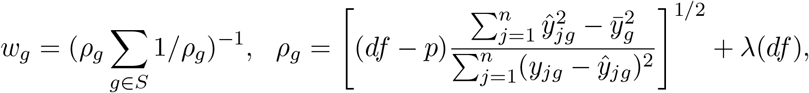

where *λ*(*df*) is a penalization that depends on *df* . The number of components *k* are selected as 1 if *df* ≤ 5, 2 if 5 < *df* ≤ 10, or 3 if *df* > 10.

#### Penalization in weights estimation

We consider a regularization *λ*(*df*) in the weights definition that depends on the degrees of freedom left in the model. When such number is low, weights estimations are imprecise and might generate artificial differences, which could lead to an even higher reduction of the underlying effective signature size.

We computed the weights average and variance for randomly selected gene sets in simulated data assuming normality and total independence between the genes. Therefore, the expected variation under gene independence due to sampling is captured and can be used in *λ*(*df*) as a way to compromise bias and precision in the weights estimations.

### 2.6 Simulations strategy

#### General procedure

We simulate data following a multivariate normal dis-tribution

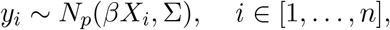

with *X*_*i*_ = 0 for *i* ≤ *n*/2 and *X*_*i*_ = 1 for *i* > *n*/2. The covariance Σ is determined by shrinking the sample correlation matrix of the metabric data (only ER+ samples) by the Identity matrix (to find a positive definite matrix). In this way we keep a similar dependence structure that can be observed in a real data set. The expected values *βX*_*i*_ are specified with regards to the scenario of testing under consideration (simulation scenarios are presented below).

We take 1,000 instances of downsampling the Metabric data to *n* = 6, 10, 20, 30. From these *n* samples, we randomly generate a factor variable that takes values 0 and 1 (*n*/2 times each) which defines the condition of interest. Moderated t-statistics are estimated for each instance of the simulation process. We use 500 rotations for approximating the p-values.

To evaluate the performance of the roastggsa scores, we compute the proportion of times (from the total 1000 instances) that the test is rejected at a significance level of 0.05.

#### Simulation scenarios

Let Ω contain all gens of the data and let *S* be a subset of Ω of size |*S*| that defines the tested gene set. We distinguish among 6 clusters of closely related genes within the gene set into the vector *C*. These groups of genes are estimated using kmeans on the original metabric data for any gene set of interest.

We identify five different scenarios to evaluate the methods under consideration: (SC0) No effect (SC1) all genes in the tested gene set have the same expected fold change which is larger than the global expected fold change; (SC2) only a group of interconnected genes in the gene set have -a common-activity in the gene set; (SC3) two groups of genes, one up-regulated and the other down-regulated, are active in the gene set; (SC4) few genes -or outliers- (in this case we set it to 3 random genes) present a much higher effect than the rest of the genes. Parameter specifications are described in eq. (2).

The SC0 is a clear consideration of a model under the null hypothesis. The SC1 and SC2 could be strategies to evaluate the power of the test, as identifying target gene sets under these two models are likely to be considered biologically relevant. The SC3 occurs rarely in public databases but its recovery might be desirable for the researcher as well. The SC4 is slightly more undesirable for its identification.

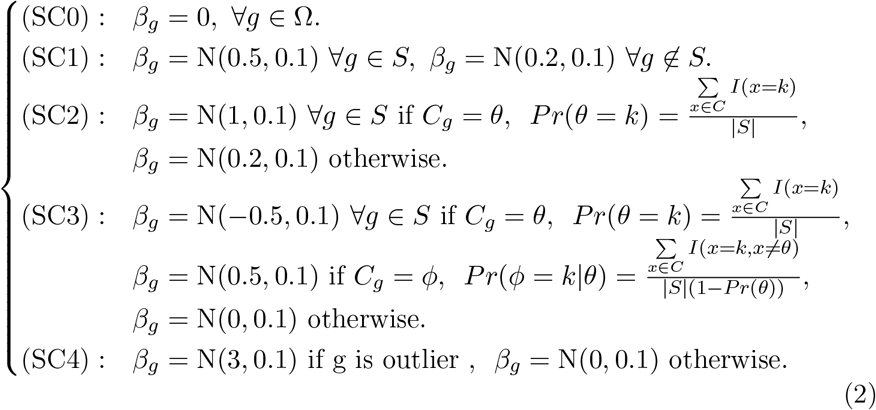

In all five scenarios we use two gene sets from the literature: genes in the (A1) TNFA signaling via NFKB pathway and genes in the (A2) interferon alpha pathway. Besides we consider an artificial case control with 31 genes that are uncorrelated (A3).

### 2.7 Measures for benchmarking results

Let [*ρ*_*i*_] be the *p*-size vector with the relevance scores of the KEGG pathways. Besides, the rank of the *i*th score in the **roastgsa** results is denoted by *I*(*i*) going from 1 (most significant) to *p* (least significant). The performance of the methods is measured using the following two measures:

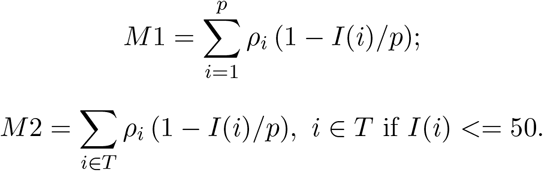

The measure M1 uses the ranks of all pathways for weighting whereas M2 only makes differences for the top 50 pathways.

## 3 Results

Specification/performance comparison of the presented **roastgsa** statistics (Table 1) is provided in Figure 2. Full results and discussion are detailed below.

**Figure 2:**
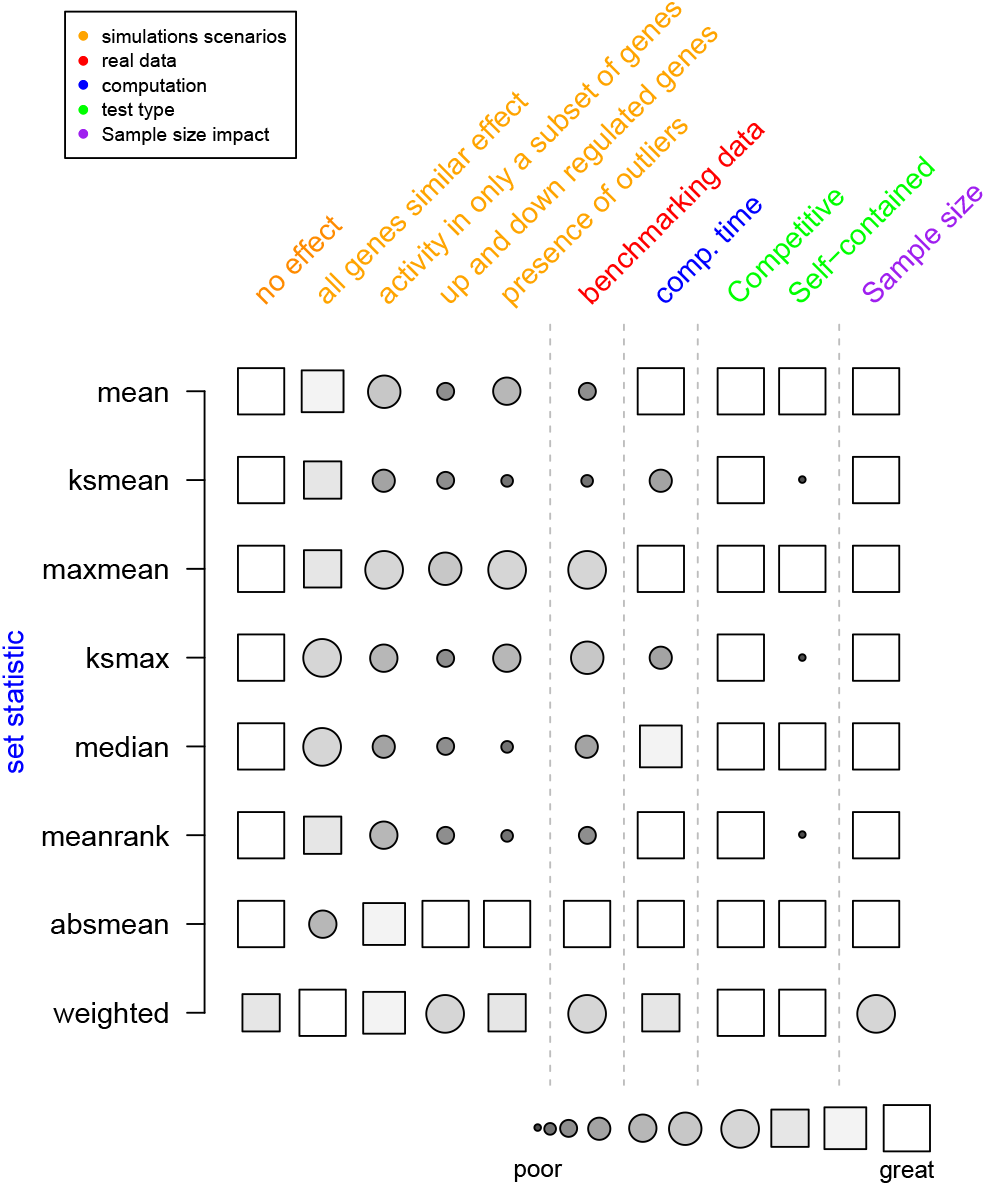
Characteristics of all presented scores: summary of performance in both simulated data and benchmarking data; Computational time; type of test; and the impact of increasing the sample size into the test. Scores go from poor performance (or not available as type of test) to excellent performance (or available as type of test), and are defined relative to all measures under study.

### 3.1 Comparison of statistics in simulated data

Five different simulation scenarios are considered as specified in Section 2.6. Empirical size (SC0) and power (SC1-SC4) are compared across roastgsa scores, complete tables presented in Supp. 2.

Weighted based scores tend to obtain more false positives than their underlying unweighted approaches in SC0. This can be due to using the testing data to also computing the weights. The impact of this bias is reduced when the sample size is sufficiently large.

On one hand, scores that are intended to capture the common activity of the pathway such as the mean, ksmean or meanrank do well in SC1 but fail to find competitive recovery rates for scenarios such as SC2, SC3 and SC4, where only few genes from the whole testing set are fixed to have a treatment effect.

On the other hand, the maxmean and absmean do not penalize for non-global activity, as it happens in more democratical scores such as the mean or meanrank, leading to the largest recovery rates for SC2, SC3 and SC4. The weighted version of the maxmean present similar powers (slightly larger) than its non-weighted counterpart, even when the testing set activity is presented in only a group of interconnected genes. Lastly, the ksmax provides poorer recovery rates than the maxmean, with the maxmean defining a much simpler statistic for interpreting the results

### 3.2 Comparison of competitive statistics in real data

The **GSEABenchmarkeR** R package is used to investigate the accuracy of the presented **roastgsa** scores in several real case studies. The package provides the relevance ranking of the KEGG pathways based on the disease under investigation (MalaCards, [16]). We consider two measures to evaluate the performance of the methods based on a weighted average of such relevance scores, one using the ordered list of all gene sets and another using only the first 50 hits [see section 2.7]. The absmean score achieves the best rates in the two measures with the maxmean being slightly better than the ksmax. In general, the weighted scores do not seem to improve the results from their underlying unweighted versions. Complete tables are presented in Supp. 3.

### 3.3 Computational complexity

In terms of computational complexity [Table 2], the absmean, mean, maxmean and mean.rank are the fastest scores to compute. The median statistic is slightly slower than the mean while the KS-based statistics take considerably larger times than the other summary statistics.

**Table 2:**
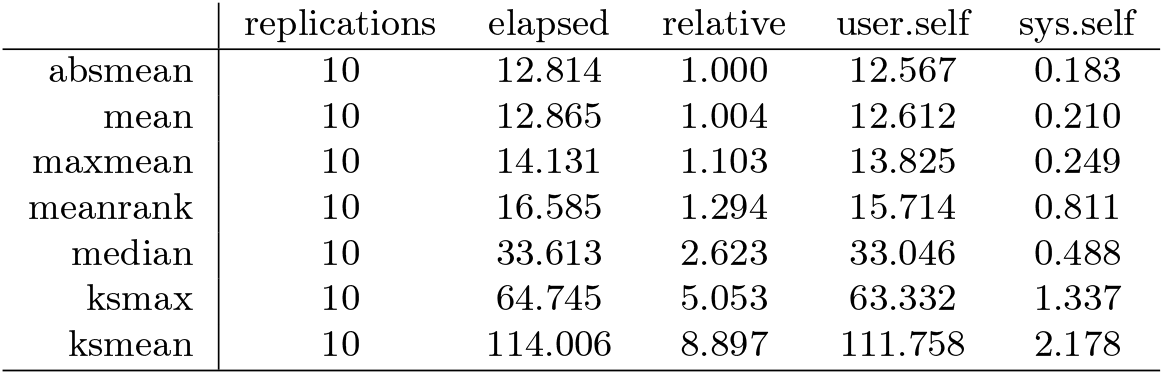
Computational time approximated in 10 replicates on 50 gene sets, 500 rotations and *N* = 50 (25 per group). Ksmax and ksmean are computationally much more intensive than the other summary statistics.

### 3.4 RoastGSA software

In the R package **roastgsa**, we propose several alternatives to visualize the results. These give the researcher some insights on the observed sample variation, gene variation and the effective signature size of every tested gene set [Figure 3].

**Figure 3:**
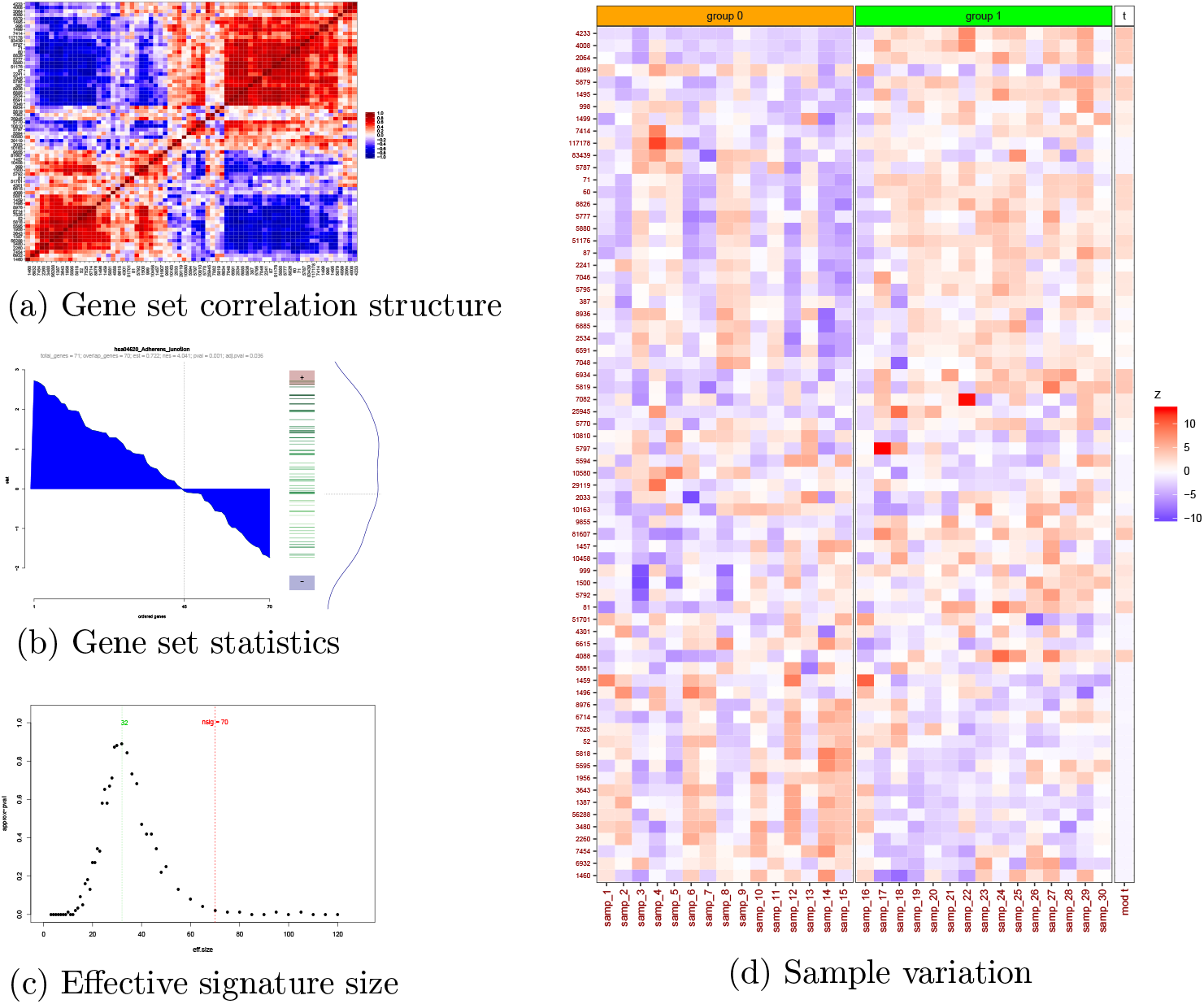
**Roastgsa** output figures: (a) heatmap of the residuals (of the linear model) correlation structure; (b) the ordered moderated t-statistics in various formats: area for up- and down- expressed genes, barcode plot for these ordered values and density; (c) Approximation of the effective signature size by comparing the variance of the rotation scores of the tested pathway with the variances of the rotation scores for sets composed by randomly selected genes of size *m*. For any effective signature size *m*, we approximate a p-value [See supplementary material 1]; (d) Normalized expression values and gene set statistics to represent the variation across samples for the gene set of interest.

## 4 Discussion

This work reviews the rotation testing approach for gene set analysis and compares the performance of the method under different enrichment score measures using both simulated and benchmarking data. The absmean (un-directional) and maxmean (directional) scores are simple measure of enrichment that have presented competitive results in all provided analyses in comparison to the more complex ksmax measure. Similarly, the mean or meanrank statistics find similar powers to ksmean, being this last much more computationally challenging. Following these empirical results, and also for the conclusive results in the work by [3], we encourage the usage of simpler measures for GSA such as the (leading method in our comparison) absmean, or the directional scores maxmean and mean.

Weighted mean and maxmean statistics are introduced as alternative to usual statistics when its expected for some genes in the pathways of interest to be heavily interconnected. These measures are found to out-power their analogous unweighted scores in the simulated data under consideration in this study, especially when all genes in the pathway play a similar role, but do not show improvements in the benchmarking data rankings.

Concerns regarding the *effective signature size* concept might represent a challenging task for researchers in the interpretation of the **roastgsa** output. A strong tendency in either side of a good part of the genes in the gene set of interest might not be corresponded with statistically relevant results, if these genes are strongly correlated. This is due to the variance of the test statistics under rotations decreasing with the effective signature size, not the signature size itself.

Published GSA figures (e.g., graphics included in Table 1) typically provide information regarding gene variation after averaging out the sample variability (i.e., taking genewise fold changes or t-statistics). We highly recommend to complement them with a graphic that shows the sample variation for the tested gene sets as well. If the dimensions are not too large, a simple heatmap, as in Figure 3d, is useful to detect those genes that are activated in the process, as quality control to detect samples that can be highly influential in the **roastgsa** analysis, and last and foremost, as a way to be honest with the total amount of data that is available for testing.

