## Supplementary material for "A comparison of rotation-based scores for gene set analysis"

Adria Caballe Mestres

January 20, 2021

### 1 Motivations for weighted summaries

#### 1.1 Results from extreme value theory

Important pathways in database collections such as Broad Hallmarks or KEGG might contain highly correlated genes, even when the effect of the known covariates is adjusted *a priori* [Figure 2a shows the correlation structure of the Interferon alpha gene set using the metabric data].

When a specific gene that is highly correlated to the rest of the gene set finds an extreme value, even under $H_0$, it is likely that many other genes in the pathway follow it with large values as well. This has been extensively studied in extreme value theory in the context of time series for asymptotic models [1], sub-asymptotic models [2] or for non-stationary processes [3]. In [4], extreme value rates for non-stationary processes that are not ordered are found to be similar to the results by [3]. We complement these theoretical results that can be found in the literature with empirical results from a case study using real data [Figure 2b].

#### 1.2 Weighted statistics and effective signature size

The weighted approach for the mean and maxmean statistics [see Table 1 in the main text] is proposed as a way to reduce the variance of the rotation scores, thus keeping larger effective signature size [Figure 3a,b]. Besides, when all genes composing the set have effects that are independent to the gene underlying correlation structure, our proposed weighting approach finds a more precise estimate of the gene set activity [Figure 3c].

*A toy example to further motivate the weighting approach.* Consider a gene set that is composed by 50 genes, forming $S_1$, that are basically explaining the same variation (in the extreme with all pair correlations equal to 1)

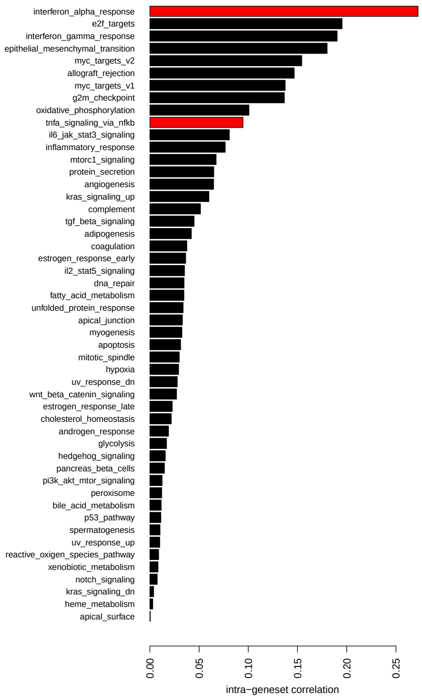


Figure 1: Intra-gene set correlation levels for all hallmarks measured by the average correlation across genes of the testing set. Interferon alpha, with the highest average correlation, and TNFA signaling via NFKB, with an intermediate correlation, are used to define correlation structures in simulated studies.

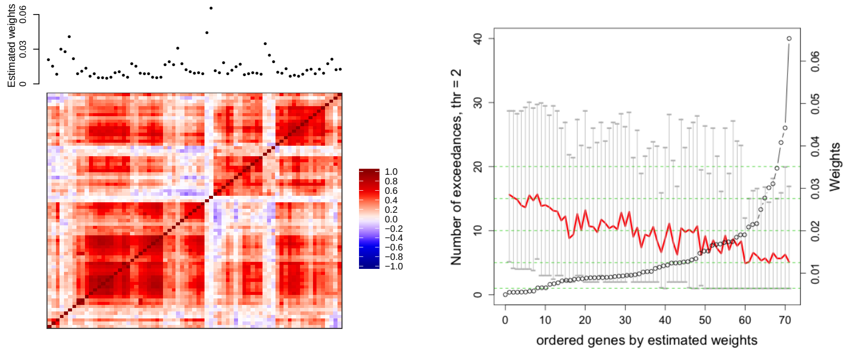


Figure 2: [Left] Heatmap with the interferon alpha gene-gene Pearson correlations (from -1 in dark blue to 1 in dark red). Estimated weights are shown above the heatmap. [Right] Number of moderated t-statistics in the interferon alpha gene set larger than 2 (in absolute value) when a given gene in the gene set had a moderated t-statistic larger than 2.

and 50 other genes, forming $S_2$, that might have similar functions in the biological process under study but that are perpendicular among each other. If we compute the mean summary statistic of this gene set, we obtain an approximated effective signature size of 4. If we re-consider the pathway so that we take all genes in $S_2$ and only 1 gene in $S_1$, the effective sample size is 51. For not such extreme case of exactly correlated genes but with strong dependency, the same sort of decrease in the effective size can occur.

### 2 Tables with recovery rates from simulation scenarios

The performance of the described methods under competitive testing is assessed using simulated data. For this purpose, we measure the proportion of times (from a total of 1000 instances) that the test is rejected at a significance level of 0.05 (from Table 1 to Table 5).

### 3 Benchmarking data results

The **GSEABenchmarkeR** R package is used to investigate the accuracy of the presented **roastgsa** scores in several real case studies. The package provides the relevance ranking of the KEGG pathways based on the disease

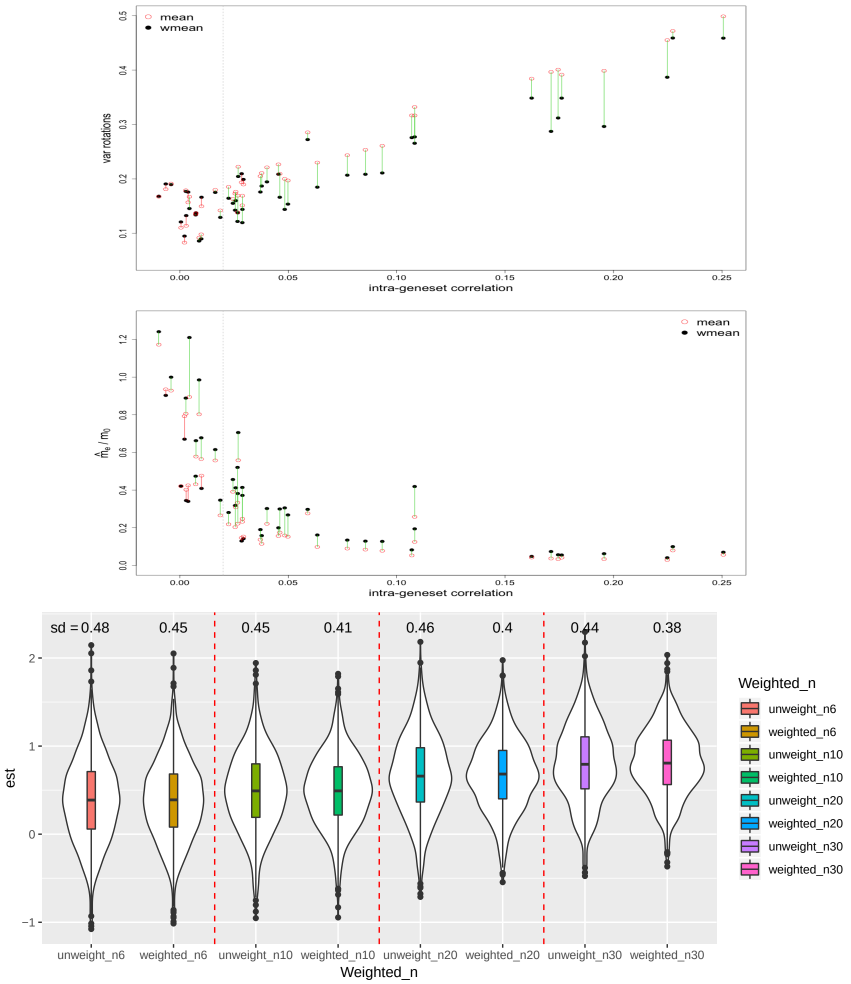


Figure 3: Weighted and unweighted mean rotation scores on Hallmarks: [top] variance of the rotation scores; [middle] ratio between estimated effective signature size and original signature size. Greater variances for the unweighted score are observed when the intra-gene set correlations is sufficiently large ($> 0.02$). This leads to a loss of effective size with respect to the weighting approach; [bottom] Self-contained mean and weighted mean enrichment scores over 2,000 instances. Constant effect of 0.3 for all genes belonging to the interferon alpha set. Weighted scores find more precise estimations than unweighted scores, especially for large $n$.

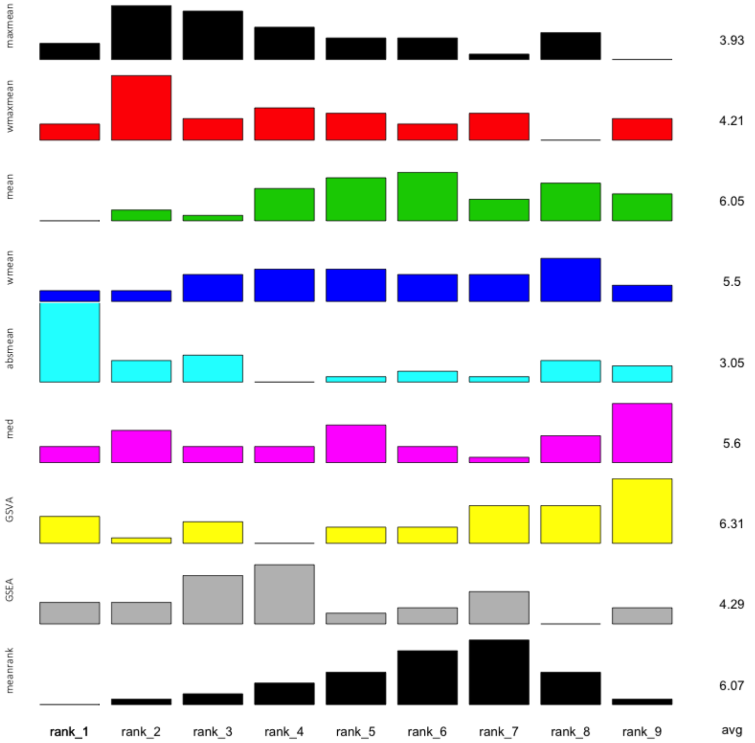


Figure 4: Barplot and average ranks resulting from measure M1 using the 42 datasets from the benchmarking package. Rank 1 is the highest rate and rank 8 is the lowest rank. Only competitive scores are used. The weighted and non-weighted maxmean scores achieve the best rates.

under investigation (MalaCards, [5]). We consider two measures to evaluate the performance of the methods based on a weighted average of such relevance scores, one using the ordered list of all gene sets and another using only the first 50 hits. The results are presented in Figure 4 (M1 measure), Figure 5 (M2 measure) and Table 6 (average ranks of M1 and M2).

[2] E. F. Eastoe and J. A. Tawn. Modelling the distribution of the clus-

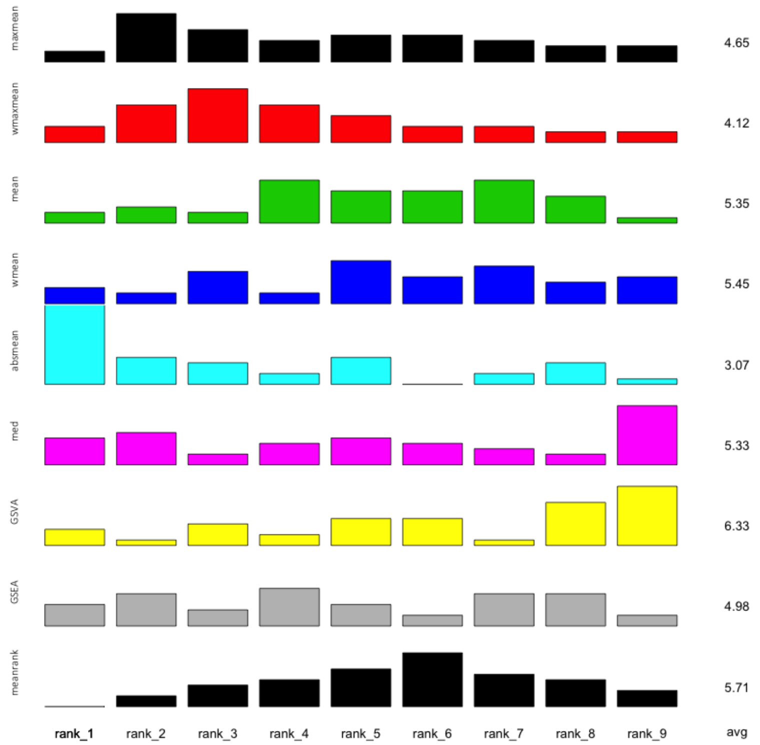


Figure 5: Average rank resulting from measures M2 using the 40 datasets from the benchmarking package. Rank 1 is the highest rate and rank 8 is the lowest rank. The maxmean score achieves the best rates in the two measures. Only competitive scores are used.

ter maxima of exceedances of subasymptotic thresholds. *Biometrika*, 99:4355, 2012.

[3] D. Aldous. Probability Approximations via the Poisson Clumping Heuristic. New York: Springer-Verlag.

[4] A. Caballe. (2018) Statistical methods for the testing and estimation of linear dependence structures on paired high-dimensional data: application to genomic data. (Doctoral dissertation), Retrieved from Edinburgh Research Archive. (http://hdl.handle.net/1842/31331).

[5] N. Rappaport, M. Twik, I. Plaschkes, R. Nudel, T. I. Stein, J. Levitt, M. Gershoni, C. P. Morrey, M. Safran, and D. Lancet. MalaCards: an amalgamated human disease compendium with diverse clinical and genetic annotation and structured search. *Nucleic Acids Research*, 45:877–887, 2017.

| n = | | 6 | 10 | 20 | 30 |
|---|---|---|---|---|---|
| | sc.mean | 45 | 53 | 50 | 47 |
| | sc.wmean | 54 | 71 | 73 | 63 |
| | sc.maxmean | 51 | 56 | 64 | 45 |
| | sc.wmaxmean | 78 | 84 | 82 | 66 |
| A1 | sc.median | 45 | 48 | 50 | 54 |
| | sc.absmean | 65 | 71 | 67 | 53 |
| | co.mean | 48 | 49 | 49 | 46 |
| | co.wmean | 60 | 66 | 76 | 61 |
| | co.maxmean | 38 | 57 | 52 | 51 |
| | co.wmaxmean | 61 | 81 | 80 | 62 |
| | co.median | 45 | 45 | 56 | 47 |
| | co.absmean | 42 | 54 | 61 | 59 |
| | meanrank | 42 | 52 | 52 | 47 |
| | ksmean | 46 | 45 | 56 | 54 |
| | ksmax | 32 | 37 | 38 | 41 |
| | sc.mean | 57 | 55 | 61 | 65 |
| | sc.wmean | 67 | 81 | 71 | 76 |
| | sc.maxmean | 59 | 64 | 51 | 64 |
| | sc.wmaxmean | 82 | 91 | 65 | 79 |
| A2 | sc.median | 54 | 66 | 56 | 61 |
| | sc.absmean | 56 | 70 | 47 | 50 |
| | co.mean | 49 | 56 | 46 | 54 |
| | co.wmean | 58 | 82 | 66 | 71 |
| | co.maxmean | 53 | 60 | 50 | 57 |
| | co.wmaxmean | 68 | 90 | 64 | 74 |
| | co.median | 52 | 67 | 49 | 55 |
| | co.absmean | 51 | 69 | 43 | 52 |
| | meanrank | 53 | 60 | 55 | 56 |
| | ksmean | 45 | 61 | 51 | 63 |
| | ksmax | 45 | 48 | 46 | 56 |
| | sc.mean | 70 | 61 | 45 | 51 |
| | sc.wmean | 61 | 53 | 37 | 55 |
| | sc.maxmean | 70 | 51 | 49 | 57 |
| | sc.wmaxmean | 79 | 56 | 50 | 64 |
| A3 | sc.median | 67 | 45 | 54 | 55 |
| | sc.absmean | 54 | 45 | 44 | 61 |
| | co.mean | 62 | 50 | 42 | 51 |
| | co.wmean | 65 | 45 | 36 | 55 |
| | co.maxmean | 63 | 45 | 59 | 62 |
| | co.wmaxmean | 76 | 50 | 47 | 64 |
| | co.median | 74 | 54 | 55 | 53 |
| | co.absmean | 51 | 46 | 44 | 66 |
| | meanrank | 66 | 48 | 49 | 53 |
| | ksmean | 54 | 44 | 41 | 54 |
| | ksmax | 55 | 49 | 37 | 57 |

Table 1: Average empirical size for all methods for $n = 6, 10, 20, 30$ iid observations ($n/2$ per group). The average intra-gene set correlation of A1, A2 and A3 are 0.05, 0.13 and 0 respectively.

| n = | | 6 | 10 | 20 | 30 |
|---|---|---|---|---|---|
| | mean | 178 | 327 | 577 | 779 |
| | wmean | 223 | 401 | 684 | **872** |
| | maxmean | 177 | 308 | 557 | 747 |
| | wmaxmean | **233** | **422** | **687** | 870 |
| A1 | median | 139 | 279 | 496 | 709 |
| | absmean | 113 | 179 | 317 | 530 |
| | meanrank | 167 | 317 | 571 | 767 |
| | ksmean | 181 | 324 | 575 | 775 |
| | ksmax | 77 | 220 | 479 | 716 |
| | mean | 86 | 126 | 231 | 334 |
| | wmean | **118** | **189** | **292** | **415** |
| | maxmean | 94 | 122 | 221 | 320 |
| | wmaxmean | 112 | 185 | 282 | 398 |
| A1 | median | 94 | 127 | 200 | 297 |
| | absmean | 75 | 121 | 175 | 259 |
| | meanrank | 81 | 130 | 236 | 329 |
| | ksmean | 91 | 132 | 236 | 322 |
| | ksmax | 66 | 129 | 212 | 303 |
| | mean | 344 | **643** | **937** | 985 |
| | wmean | **348** | 607 | 926 | **986** |
| | maxmean | 312 | 589 | 910 | 979 |
| | wmaxmean | 304 | 580 | 900 | 980 |
| A1 | median | 210 | 410 | 793 | 911 |
| | absmean | 114 | 175 | 431 | 618 |
| | meanrank | 335 | 594 | 913 | 984 |
| | ksmean | 294 | 564 | 906 | 972 |
| | ksmax | 129 | 424 | 859 | 966 |

Table 2: Proportion ($\times$ 1000) of tests that have been rejected with a rejection levels of 0.05. Scenario 1: same effect for all genes in the tested geneset, $n = 6, 10, 20, 30$ iid observations (n/2 for each group). Only competitive test statistics are shown. The average intra-gene set correlation of A1, A2 and A3 are 0.05, 0.13 and 0 respectively.

| n = | | 6 | 10 | 20 | 30 |
|---|---|---|---|---|---|
| | mean | 74 | 91 | 170 | 268 |
| | wmean | 95 | 125 | 227 | 334 |
| | maxmean | 88 | 103 | 221 | 334 |
| | wmaxmean | **137** | 179 | 307 | 433 |
| A1 | median | 65 | 75 | 120 | 173 |
| | absmean | 115 | **197** | **407** | **646** |
| | meanrank | 74 | 74 | 129 | 171 |
| | ksmean | 60 | 79 | 126 | 180 |
| | ksmax | 37 | 59 | 127 | 223 |
| | mean | 74 | 69 | 130 | 144 |
| | wmean | 96 | 105 | 151 | 164 |
| | maxmean | 82 | 76 | 145 | 182 |
| | wmaxmean | **110** | 118 | 182 | 206 |
| A2 | median | 71 | 59 | 107 | 116 |
| | absmean | 102 | **124** | **236** | **353** |
| | meanrank | 65 | 49 | 101 | 97 |
| | ksmean | 63 | 50 | 99 | 102 |
| | ksmax | 55 | 61 | 107 | 144 |
| | mean | 168 | 284 | 499 | 680 |
| | wmean | 153 | 270 | 491 | 668 |
| | maxmean | **175** | 352 | **593** | **764** |
| | wmaxmean | **175** | **355** | 589 | 757 |
| A3 | median | 106 | 161 | 234 | 294 |
| | absmean | 127 | 270 | 525 | 727 |
| | meanrank | 117 | 189 | 296 | 404 |
| | ksmean | 117 | 153 | 307 | 431 |
| | ksmax | 64 | 158 | 451 | 685 |

Table 3: Proportion ($\times$ 1000) of tests that have been rejected with a rejection levels of 0.05. Scenario 2: only a group of interconnected genes in the geneset have -a common- activity in the geneset, $n = 6, 10, 20, 30$ iid observations (n/2 for each group). Only competitive test statistics are shown. The average intra-gene set correlation of A1, A2 and A3 are 0.05, 0.13 and 0 respectively.

| n = | | 6 | 10 | 20 | 30 |
|---|---|---|---|---|---|
| | mean | 59 | 56 | 63 | 67 |
| | wmean | 79 | 73 | 100 | 104 |
| | maxmean | 80 | 78 | 114 | 173 |
| | wmaxmean | **114** | 127 | 176 | 267 |
| A1 | median | 62 | 46 | 55 | 64 |
| | absmean | 112 | **184** | **378** | **590** |
| | meanrank | 51 | 42 | 46 | 47 |
| | ksmean | 58 | 45 | 44 | 41 |
| | ksmax | 44 | 43 | 63 | 73 |
| | mean | 67 | 43 | 48 | 60 |
| | wmean | 86 | 67 | 64 | 77 |
| | maxmean | 63 | 53 | 77 | 102 |
| | wmaxmean | **98** | 88 | 102 | 127 |
| A1 | median | 58 | 42 | 48 | 62 |
| | absmean | 82 | **101** | **175** | **259** |
| | meanrank | 58 | 32 | 36 | 40 |
| | ksmean | 56 | 34 | 39 | 38 |
| | ksmax | 56 | 41 | 51 | 49 |
| | mean | 70 | 85 | 96 | 101 |
| | wmean | 61 | 74 | 78 | 97 |
| | maxmean | 126 | 151 | 335 | 462 |
| | wmaxmean | **132** | 152 | 343 | 457 |
| A1 | median | 63 | 82 | 94 | 109 |
| | absmean | 129 | **194** | **504** | **690** |
| | meanrank | 51 | 63 | 61 | 64 |
| | ksmean | 46 | 55 | 49 | 42 |
| | ksmax | 68 | 88 | 117 | 144 |

Table 4: Proportion ($\times$ 1000) of tests that have been rejected with a rejection levels of 0.05. Scenario 3: two groups of genes, one up-regulated and the other down-regulated, are active in the geneset, $n = 6, 10, 20, 30$ iid observations (n/2 for each group). Only competitive test statistics are shown. The average intra-gene set correlation of A1, A2 and A3 are 0.05, 0.13 and 0 respectively.

| n = | | 6 | 10 | 20 | 30 |
|---|---|---|---|---|---|
| A1 | mean | 50 | 71 | 55 | 82 |
| | wmean | 61 | 97 | 80 | 110 |
| | maxmean | 80 | 94 | 88 | 126 |
| | wmaxmean | **113** | 152 | 152 | 181 |
| | median | 47 | 50 | 54 | 56 |
| | absmean | 107 | **155** | **238** | **305** |
| | meanrank | 44 | 61 | 43 | 47 |
| | ksmean | 37 | 59 | 41 | 43 |
| | ksmax | 41 | 62 | 42 | 63 |
| A2 | mean | 58 | 76 | 90 | 86 |
| | wmean | 80 | 106 | 114 | 126 |
| | maxmean | 81 | 97 | 120 | 134 |
| | wmaxmean | 110 | 141 | 174 | 197 |
| | median | 57 | 53 | 53 | 50 |
| | absmean | **141** | **190** | **326** | **465** |
| | meanrank | 49 | 54 | 47 | 46 |
| | ksmean | 51 | 43 | 32 | 38 |
| | ksmax | 52 | 60 | 63 | 65 |
| A3 | mean | 472 | 715 | 934 | 979 |
| | wmean | 434 | 693 | 925 | 970 |
| | maxmean | **690** | **949** | **1000** | **1000** |
| | wmaxmean | 675 | 932 | **1000** | **1000** |
| | median | 59 | 80 | 75 | 87 |
| | absmean | 615 | 928 | **1000** | **1000** |
| | meanrank | 71 | 94 | 108 | 111 |
| | ksmean | 86 | 110 | 135 | 115 |
| | ksmax | 514 | 796 | 985 | 999 |

Table 5: Proportion ($\times$ 1000) of tests that have been rejected with a rejection levels of 0.05. Scenario 4: few genes -or outliers- present a much higher effect than the rest of the genes, $n = 6, 10, 20, 30$ iid observations (n/2 for each group). Only competitive test statistics are shown. The average intra-gene set correlation of A1, A2 and A3 are 0.05, 0.13 and 0 respectively.

| | maxmean | wmaxmean | mean | wmean | absmean | med | ksmean | ksmax | meanrank |
|---|---|---|---|---|---|---|---|---|---|
| M1 | 3.93 | 4.21 | 6.05 | 5.50 | **3.05** | 5.60 | 6.31 | 4.29 | 6.07 |
| M2 | 4.66 | 4.12 | 5.35 | 5.45 | **3.07** | 5.33 | 6.33 | 4.98 | 5.71 |

Table 6: Average rank resulting from measures M1 and M2 using the 40 datasets from the benchmarking package. Rank 1 is the highest rate and rank 9 is the lowest rank. The absmean score achieves the best rates in the two measures followed by the maxmean approach. Only competitive scores are used.
